# Whole Genome Sequencing of Antimicrobial-Resistant *Shigella sonnei* Associated with Infection Acquired from International and Domestic Settings Reveals Gene Allelic Variants that Predict Global Lineages

**DOI:** 10.1101/361196

**Authors:** Rebecca L. Abelman, Nkuchia M. M’ikanatha, Edward G. Dudley

**Affiliations:** Department of Food Science, The Pennsylvania State University, University Park, Pennsylvania, USA; *E. coli* Reference Center, The Pennsylvania State University, University Park, Pennsylvania, USA; Pennsylvania Department of Health, Harrisburg, Pennsylvania, USA

## Abstract

*Shigella spp*. are a major cause of gastroenteritis worldwide, and *S. sonnei* is the most common species isolated within the United States. Recently, advancements in technology have made whole genome sequencing (WGS) readily available, and as such, laboratories are moving to implement WGS in outbreak analysis, surveillance, and antimicrobial resistance (AMR) monitoring of major foodborne pathogens. Accordingly, our study examined a collection of 22 antimicrobial resistant *S. sonnei* isolates from patients who either acquired the infections within the United States or when travelling to international locations between 2009 to 2014. We applied WGS to investigate both the relatedness of these isolates and the genetic determinants of AMR to address the phenotypic differences seen in previous observations. We analyzed the phylogeny of these strains and observed segmentation based on the previously described Global Lineages of *S. sonnei*. Following these results, 17 gene sequences with lineage specific single nucleotide polymorphisms (SNPs) were identified and developed into a lineage prediction test to determine the Global Lineage of uncharacterized *S. sonnei*, which accurately predicted phylogenetic segmentation and additionally showed specificity for *S. sonnei* genomes (97% accuracy, 38/39 genomes). Lastly, to determine differences between either the international or domestic isolates or between the Global Lineages, the AMR determinants were identified. We found a variety of AMR determinants within the genomes, and while the international and domestic *S. sonnei* carried similar resistance determinants, differences between Global Lineages were observed.

## Introduction

*Shigella spp*. are agents of bacillary gastrointestinal illness responsible for an estimated 80 to 165 million cases worldwide (1, 2); consequently, they rank as the fourth most common cause of bacterial foodborne illness in the United States (3). There are four species of *Shigella*; in the United States and other developed countries, *Shigella sonnei* accounts for over 80% of all shigellosis infections (4, 5). Research on *S. sonnei* has increased to investigate both the previously under-described nature of this species and the shift in population within developed regions from *S. flexneri* dominance to *S. sonnei* in recent history (6). Though many theories have been suggested and the root cause of the shift from *S. flexneri* to *S. sonnei* still remains unknown, additional research on *S. sonnei* is vital.

Characterization of *S. sonnei* has traditionally depended on pulsed-field gel electrophoresis (PFGE) (7), but whole genome sequencing (WGS) has recently gained popularity due to its higher resolution, increased availability, and lower costs. Accordingly, government and state laboratories alike are updating characterization and surveillance techniques to utilize WGS. One area lacking in observation is the phylogenomics of *S. sonnei*, specifically in the *S. sonnei* isolated within the United States and how it relates to those found internationally. Holt et al. described a specific clustering pattern within *S. sonnei* phylogeny known as the Global Lineage hypothesis, which showed that *S. sonnei* segment into 4 main clusters on a phylogenetic tree made using SNP variant analysis (8). However, this study did not include any *S. sonnei* from the United States. This hypothesis has only been tested once on *S. sonnei* strains found in the United States, when a group from California used WGS to investigate two separate outbreaks of *S. sonnei*; they found that most isolates fell into Lineage III (9). However, the scope of this analysis was narrow, as it was only based around two outbreaks within California, so studies with a wider range of isolates may yield additional beneficial data.

As with many other foodborne pathogens, antimicrobial resistance (AMR) in *S. sonnei* is (NARMS) in 2014, 2.4% of all *Shigella* isolated from humans in the United States were resistant on the rise. In a recent report by the National Antimicrobial Resistance Monitoring System to ciprofloxacin and 4.7% were resistant to azithromycin, the first-line antibiotics typically used to treat shigellosis (10). Additionally, the report showed that over 40% of all *Shigella* isolated were resistant to three or more classes of antibiotics. Traditionally, testing and surveillance of antibiotic susceptibility is performed via standardized *in vitro* methods (11), however detection of AMR determinants directly from the genome shows promise as a possible replacement for *in vitro* testing methodology. Comparisons between *in vitro* testing and genetic testing for AMR have been investigated in *Escherichia coli* and other gram-negative bacteria, and these studies have found greater than 97% correlation between both methods (12–15). To our knowledge, only one study has investigated the correlation of phenotypic *S. sonnei* antimicrobial susceptibility results to genome antimicrobial analysis of the same strains, and they found similar correlation results (9). Additional research is needed to gauge the effectiveness of AMR detection via genome analysis when compared to traditional *in vitro* testing, specifically on a wider range of *S. sonnei* isolates.

Previously, the Pennsylvania Department of Health performed a wide scale analysis on a Health collected travel history from these patients and grouped the *Shigella* as international isolates and domestic isolates. Using these *Shigella*, they examined the correspondence of geographic origin and PFGE to antimicrobial susceptibility, and they found that international isolates carried more antibiotic resistance (16). In response to these results, 22 *S. sonnei* from the same collection were chosen for WGS to examine in greater detail; specifically, we performed a phylogenetic analysis to view relatedness between the isolates and examined the AMR determinants within the genomes. Based on the previous results generated by Li et al. (16), we expected to observe higher rates of AMR encoded by a greater number of genes in the international isolates, and we also expected phylogenetic segmentation based on geographic origin. Yet, we observed segmentation collection of *Shigella* isolated from Pennsylvania patients (16). The Pennsylvania Department of based on the Global Lineages, so genes that could be used to predict phylogenetic classification were identified and tested for validity.

## Materials and Methods

### Bacterial strains and growth conditions

The *S. sonnei* isolates sequenced in this study (Table 1) were previously analyzed by the Pennsylvania Department of Health as part of AMR surveillance of enteric bacteria (16). After receiving the bacterial isolates from the public health laboratory, they were promptly inoculated from the shipping stabs to sterile Lysogeny Broth (LB) agar plates and grown overnight at 37°C.

**Table 1:** *S. sonnei* isolates sequenced in this study

| PSU-ID<sup>a</sup> | Travel History<sup>b</sup> | Year of Isolation | Antimicrobial Resistance Profile<sup>c</sup> |
| --- | --- | --- | --- |
| SS-2 | None | 2011 | STR, AMP, AMC, FOX, TET |
| SS-3 | None | 2013 | GEN, STR, SXT, SMX, TET, AZM |
| SS-4 | None | 2013 | STR, SXT, SMX, TET |
| SS-5 | None | 2014 | STR, AMP, SXT, SMX, TET, AZM |
| SS-21 | India | 2009 | STR, SXT, NAL, SMX, TET |
| SS-23 | Nepal | 2009 | STR, SXT, CIP, NAL, SMX, TET |
| SS-24 | India | 2010 | STR, SXT, CIP, NAL, SMX, TET |
| SS-26 | Jamaica | 2010 | STR, SXT, NAL, SMX, TET |
| SS-27 | None | 2010 | STR, AMP, SXT, SMX, TET |
| SS-28 | None | 2011 | STR, AMP, AMC, FOX |
| SS-29 | None | 2011 | STR, AMP, AMC, FOX |
| SS-30 | Peru | 2012 | AMP, AMC, SXT, CHL, SMX, TET |
| **SS-31** | Peru | 2012 | AMP, AMC, SXT, CHL, SMX, TET |
| **SS-32** | None | 2012 | STR, AMP, SXT, SMX, TET |
| **SS-35** | India | 2012 | STR, SXT, CIP, NAL, SMX, TET |
| **SS-36** | None | 2012 | STR, SXT, CIP, NAL, SMX, TET |
| **SS-37** | Haiti | 2013 | STR, AMP, SXT, SMX, TET |
| **SS-38** | Dominican Republic | 2013 | STR, AMP, SXT, SMX, TET |
| **SS-39** | None | 2013 | STR, SXT, CIP, NAL SMX, TET |
| **SS-40** | None | 2013 | STR, AMP, SXT, SMX, TET |
| **SS-42** | Dominican Republic | 2014 | STR, SXT, NAL, SMX, TET |
| **SS-43** | Mexico | 2014 | STR, SXT, NAL, SMX, TET |

<sup>a</sup>Specimen identifications in the form of PSU-IDs were assigned to each *S. sonnei* isolate received from the Pennsylvania Department of Health.
<sup>b</sup>Travel history describes where the patient is believed to have acquired the *S. sonnei* infection. “None” indicates that the patient did not report travelling to an international location, thus the *S. sonnei* was acquired domestically.
<sup>c</sup>Antimicrobials listed within this chart were tested and found resistant based on previous data (16). Abbreviations are as follows: STR for Streptomycin, GEN for Gentamicin, AMP for Ampicillin, AMC for Amoxicillin and Clavulanic Acid, FOX for Cefoxitin, SXT for Trimethoprim-sulfamethoxazole, SMX for Sulfamethoxazole, TET for Tetracycline, CHL for Chloramphenicol, CIP for Ciprofloxacin, NAL for Nalidixic Acid, AZM for Azithromycin.

### DNA extraction, library preparation, and whole genome sequencing

For DNA extraction, a pure colony from each overnight agar culture was inoculated into 3 mL of LB for overnight growth at 37°C in a shaking incubator at 300 RPM. DNA extraction was performed using a Wizard Genomic DNA kit (Promega, Madison, WI), following the manufacturer’s instructions. DNA concentration was quantified on a Qubit 2.0 fluorometer using the Qubit dsDNA BR Assay kit (Thermo-Fisher, Waltham, MA) and DNA purity was checked using a A260/A280 purity ratio. Target DNA concentration was at or greater than 10 ng/μL and target purity ratio was greater than 1.8 and less than 2.2. The DNA of each strain was then diluted to a concentration of 0.2 ng/μL and prepared into sequencing libraries using the Nextera XT DNA Library Preparation Kit (Illumina, Inc., San Diego, CA) per manufacturer’s instructions. An Illumina MiSeq was used to sequence the isolates, using 250 bp paired-end read length sequencing chemistry.

### Read assembly and quality control

Sequencing read quality was initially determined using FastQC 0.11.5 (17), followed by assembly using the SPAdes Genome Assembler Version 3.10.0 (18) with default parameters. Assembled genomes were then run in QUAST 4.5 (19) to determine the number of contigs, the N50 score, and the total length of the assembled genome. Lastly, read coverage was calculated by aligning the reads of each strain to a reference *S. sonnei* genome using the Burrow-Wheeler Aligner 0.7.15 (20) and SAMtools 0.1.18 (21) to create a BAM file. Next, the SAMtools depth function was used to calculate the average read coverage. Once collected, the generated parameters were compared to the values used for sequencing of *Escherichia coli* by the Centers for Disease Control and Prevention (22). All quality control values are reported in Table S1, and all sequences were considered acceptable.

### Antimicrobial resistance gene identification

Assembled genomes were screened using BLAST+ (23) against the Bacterial Antimicrobial Resistance Reference Gene Database (BARRGD) to identify AMR genes (BioProject number PRJNA313047, accessed May 2017). Genes with high nucleotide percent identity (>99% identity) and high query coverage (>99% coverage) were marked as present within the genome, and the genomes were next screened using the ResFinder 3.0 database (24), using default parameters (>90% nucleotide percent identity and 60% query coverage). Only genes identified by both BARRGD and ResFinder were used for comparisons of phenotypic results. Additionally, ResFinder was used to identify known chromosomal mutations that result in AMR phenotypes.

### Phylogenetic analysis

The Single Nucleotide Variant Phylogenomics (SNVPhyl) pipeline was used to perform SNP calling between the *S. sonnei* isolates and to construct the resulting phylogenetic tree (25). All parameters were kept at default settings, and the minimum coverage was set to 10, the minimum mean mapping quality score was set to 30, and the SNV abundance ratio was set to 0.75 as suggested. *S. sonnei* Ss046 (NCBI Accession Number NC_007384.1/CP_000038.1) was used as the reference genome for all SNVPhyl runs. The output of SNVPhyl included a maximum likelihood phylogenetic tree generated by PhyML, a SNP distance matrix table, and a SNP table with all called SNPs and their locations compared to the reference genome. To calculate bootstrap values, the phylogenetic tree was reanalyzed by PhyML 3.0 (26). Some phylogenetic trees were visualized using Microreact (27).

### Identification of presumptive lineage specific alleles

Putative lineage specific alleles were identified by using the SNVPhyl generated SNP Table. The SNP Table was reviewed manually, and only SNPs considered “Valid” by the SNVPhyl workflow were considered in this analysis. Locations of lineage specific SNPs were determined by finding SNPs that occurred only in Lineage III strains or only in Lineage II strains. These SNP positions were noted and then compared to the annotation of the reference genome, to locate genes that carry these lineage specific SNPs. The full sequences of each gene that had a lineage specific SNP were extracted from the *S. sonnei* Ss046 genome and saved for confirmation testing.

### Confirmation testing and usage of lineage specific alleles

Once the sequences of the presumptive lineage specific alleles were collected, 13 additional *S. sonnei* genomes in the form of short reads were pulled from the NCBI Sequence Read Archive (SRA) using the SRA Toolkit Version 2.8.1 (28). The reads were assembled and the quality of the reads were checked using the methods described above. Additionally, SNVPhyl was used to perform SNP calling on these *S. sonnei* and create a phylogenetic tree and depict the global lineages. The assembled genomes were then aligned to the lineage specific alleles using BLAST+, and sequences with consistency between *S. sonnei* of the same global lineage were considered confirmed lineage specific (i.e. if all Lineage II *S. sonnei* had the allele at 100% nucleotide identity and all Lineage III *S. sonnei* had the same allele at 99% nucleotide identity, the gene allelic variant was considered a Lineage II specific allele). This process removed presumptive lineage specific alleles that were inconsistent or possible artifacts of the initial SNP calling process. These confirmed lineage specific alleles can then be used to predict the Global Lineage of *S. sonnei* by aligning them via BLAST to the assembled genome of question. If the *S. sonnei* aligns to Lineage II alleles at 100% nucleotide identity, it is a predicted Lineage II isolate, and the inverse is true for Lineage III *S. sonnei*. Furthermore, lineage prediction could be performed by locating the specified SNPs within the alleles; Lineage III isolates all carry the same listed SNP for the Lineage II alleles while the Lineage II isolates have the specified SNPs in the Lineage III alleles. This prediction method also distinguished *S. sonnei* from other *Shigella* species and *E. coli*, as isolates that are not *S. sonnei* will have less than 100% nucleotide identity to all confirmed lineage specific alleles.

### Accession numbers

Isolates sequenced in this study and corresponding accession numbers can be found in Table S2. They were uploaded to the National Center for Biotechnology Information (NCBI) under BioProject Number PRJNA273284. Other sequences utilized in this study can be found in Table S3.

## Results

### Phylogenetic analysis of *S. sonnei* Collection

SNP calling was performed and a maximum-likelihood tree was generated to visualize the relatedness of these *S. sonnei*; clustering based on geographic origin of the strains was expected to be the main driver of phylogenetic segmentation. The *S. sonnei* separated into two distinct clusters, separated by > 850 SNPs, with the isolates SS-32, SS-37, SS-38, and SS-40 segmenting distinctly from the remaining eighteen isolates (Fig. 1). Additional branching occurred within the two main clusters; in some cases, like with SS-30 and SS-31, highly related strains formed separate smaller clusters. However, overall clustering occurred seemingly without the influence of geographic origin, as both of the main clusters contained international and domestic isolates. To further investigate the phylogenetic segmentation, 17 isolates from the Holt collection (29) were included in our analysis approach to test whether the Global Lineages could explain the observed clustering (Fig. 2). When analyzed with representative strains from the Holt *S. sonnei* collection, the Pennsylvania *S. sonnei* sequences segmented into Lineage II or III, mirroring the 2 main clusters in Fig. 1A, with 4 isolates aligning with strains in Lineage II and the other 18 isolates falling into Lineage III. No strains clustered into Lineage I or IV.

**Figure 1:**
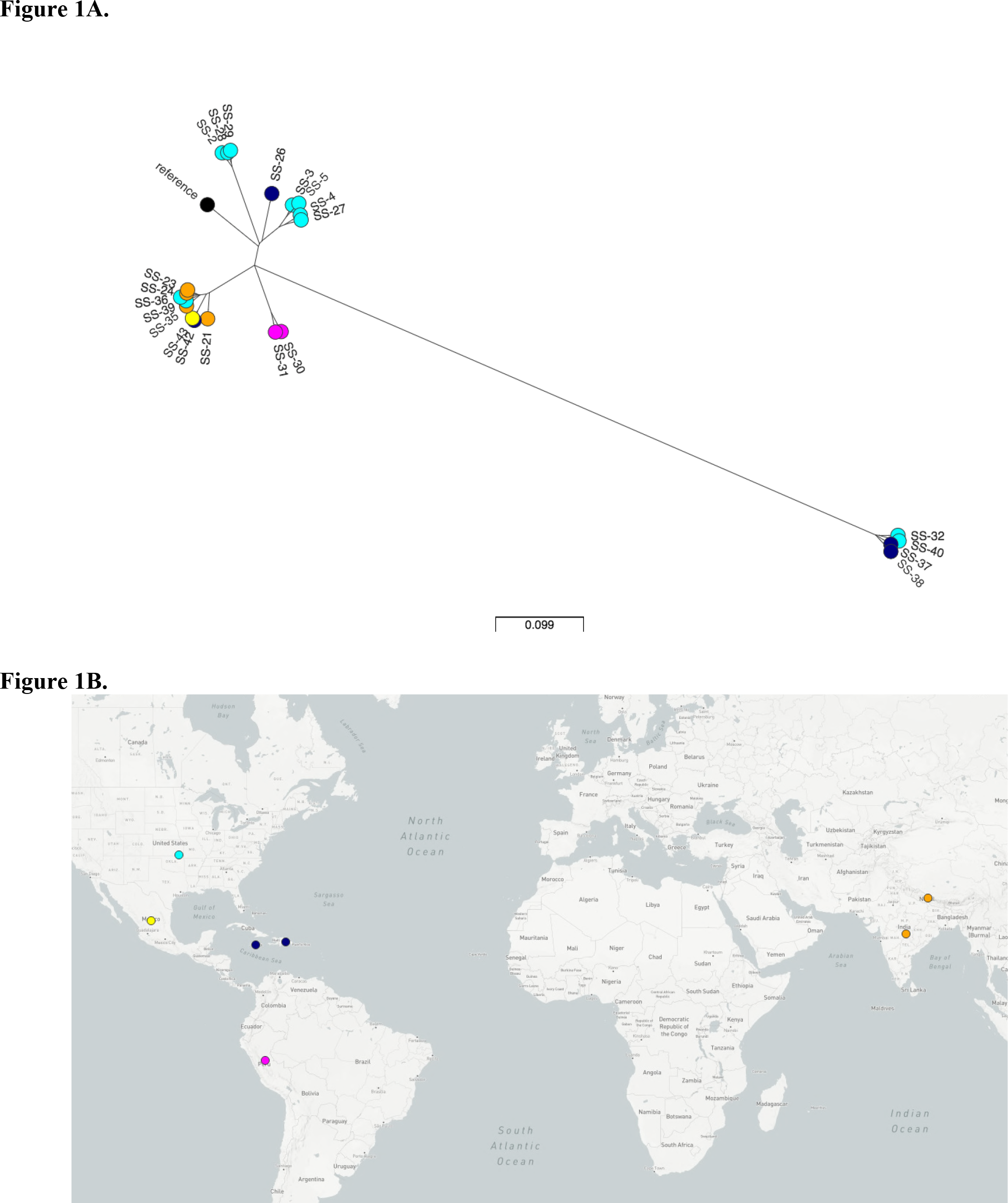
*S. sonnei* Phylogenetic Tree and World Map via Microreact. *S. sonnei* Phylogenetic Tree (Figure 1A) and World Map (Figure 1B) generated via the Microreact Program. The colored circles of the tree correspond with the matching colored circles on the map. Colors were designated by region, and the location of the points do not correspond with the exact geographic location within the country, but rather the country as a whole. An interactive version of this output can be found at https://microreact.org/project/rJxfWLQdb.

**Figure 2:**
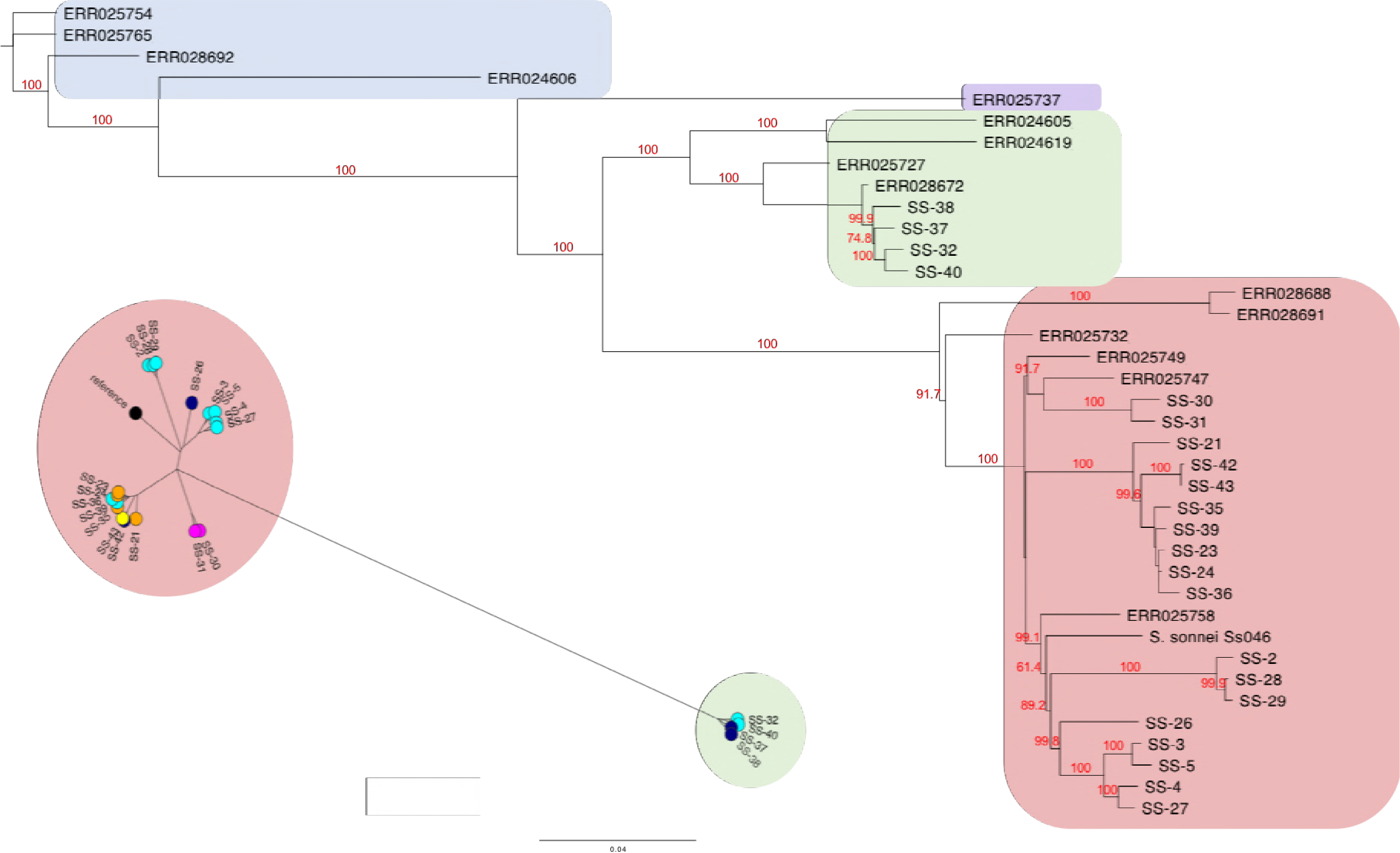
Global lineage analysis of *S. sonnei* collection. A GTR gamma maximum likelihood phylogenetic tree was generated by SNVPhyl to compare the Global Lineage strains to the *S. sonnei* collection, and bootstraps were generated by reanalyzing the tree using PhyML. The blue highlighted region contains isolates in Linage I, the purple contains Lineage IV, the green contains Lineage II, and the red contains Lineage III. Figure 2 also features Figure 1A with highlighted lineage regions mirroring the pattern seen.

### Global Lineage prediction gene allelic variants

Thirteen Lineage III specific alleles and four Lineage II alleles sequences that carried lineage-specific SNPs were identified from the 22 *S. sonnei* isolates (Table 2). To test the validity of using these sequences as a prediction tool, an additional 39 *S. sonnei* genomes were downloaded from NCBI’s SRA, and the 17 confirmed lineage specific alleles were then applied to predict the Global Lineage of each isolate. Lineages were predicted by comparing the identity of the Lineage II specific alleles versus the Lineage III specific alleles (Table 3). In rare cases, Lineage III *S. sonnei* isolates would carry all but one lineage specific sequences of the 13 Lineage III group at 100% nucleotide identity. There was no consistency to this discrepancy, but these isolates were still predicted as Lineage III based on the majority of identical sequences. When Lineage III isolates did not carry a Lineage III allele at full identity, it typically occurred due to an acquired SNP that was distinct from the SNP carried by Lineage II isolates. Predictions by the allele sequences were verified by performing the full phylogenetic analysis and locating the isolates within the lineages on the phylogenetic tree (Fig. 3).

**Figure 3:**
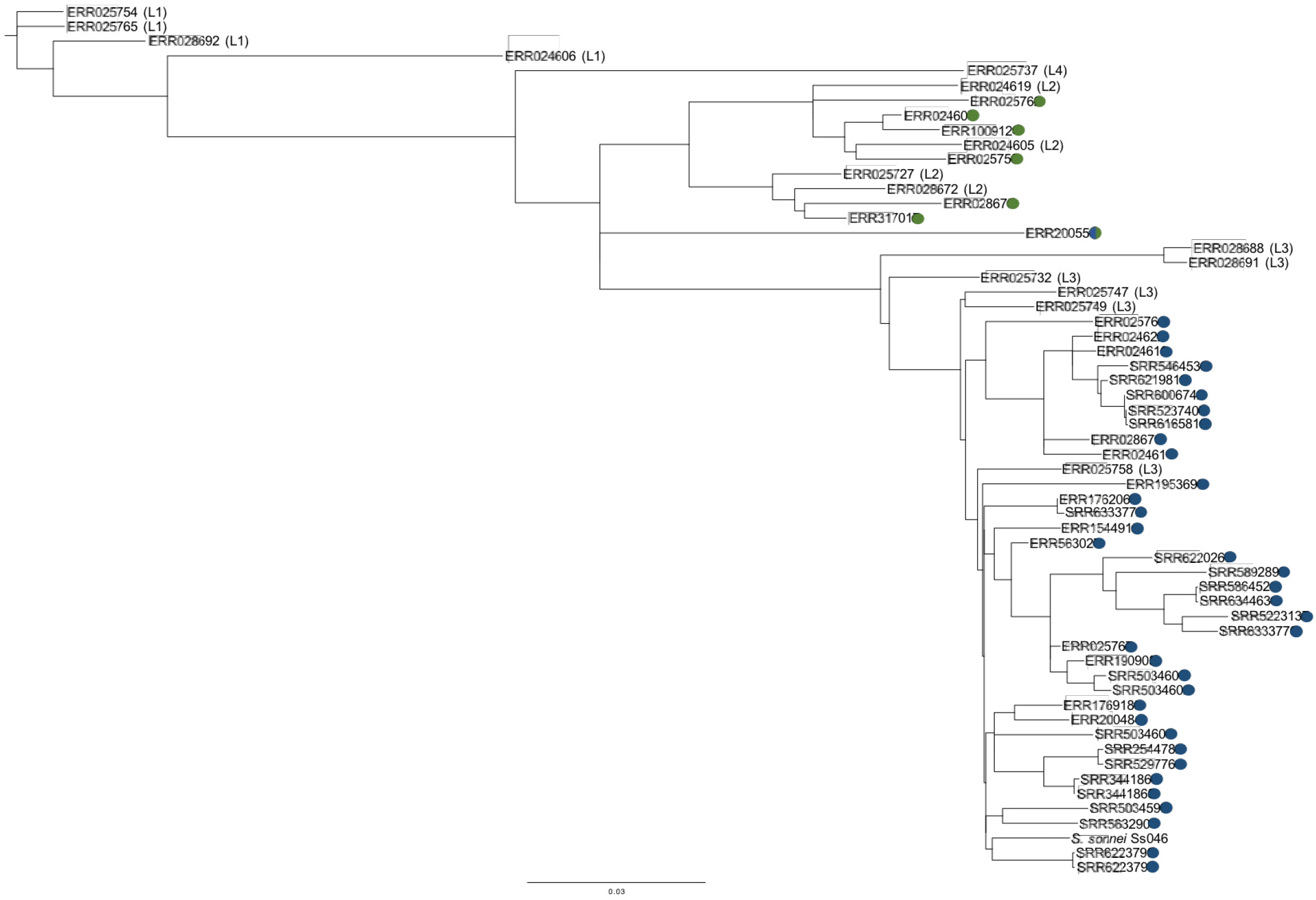
Confirmation of Global Lineage Prediction. This SNVPhyl-generated phylogenetic tree serves as the confirmatory Global Lineage identifier for the additional *S. sonnei* isolates used to test the lineage specific alleles. Sequencing reads for each isolate were pulled from the NCBI SRA, then inputted into SNVPhyl along with the 17 Global Lineage *S. sonnei* strains from the Holt et al. collection to perform SNP calling. The resulting tree was compared to the global lineage predictions (Table 3), and the colored dots represent what the initial lineage prediction was: blue dots for predicted Lineage III and green dots for Lineage II. *S. sonnei* ERR200550 received a special designation of a half-blue/half-green dot, as it carried lineage specific alleles for both Lineage II and III and was considered “Not Predictable.”

**Table 2:**
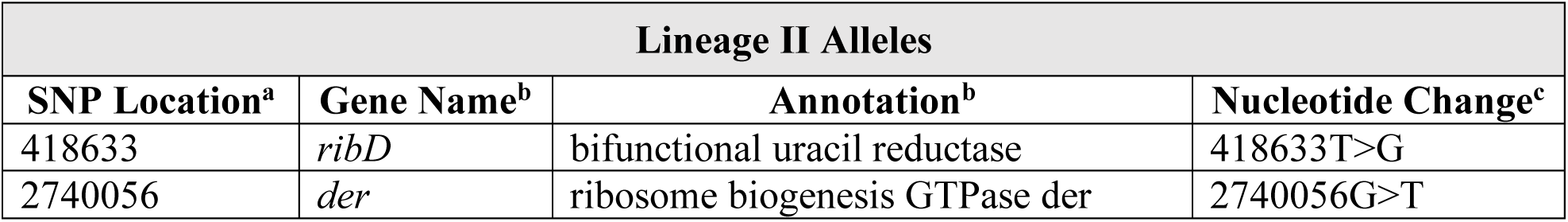
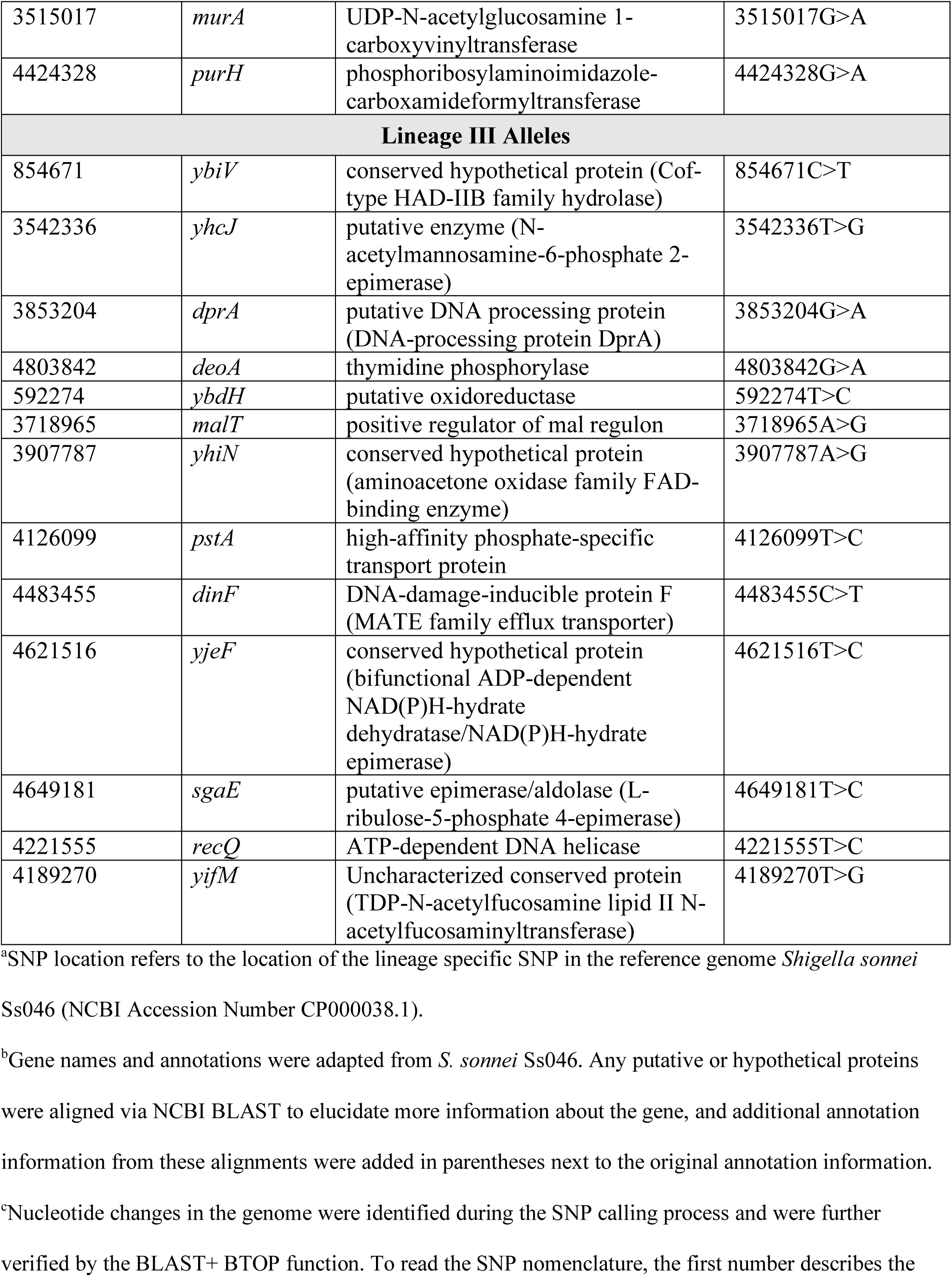
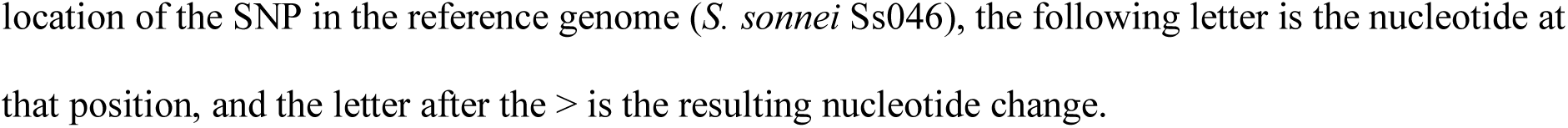
Confirmed Lineage Specific Alleles Sequences

Of the 39 additional *S. sonnei* sequences tested by this prediction assay, only one of them was unable to be characterized. This isolate, ERR200550, carried various Lineage II and III equences at 100% nucleotide identity, and it did not follow the same identity patterns as the other Lineage II or Lineage III isolates. Upon using a full phylogenetic analysis, it appeared that this strain did not fall in any of the previously described lineages, but instead fell into a branched distinctly off the main brain separating Lineage II and Lineage III. In addition, the lineage specific alleles showed high specificity to *S. sonnei* when compared to other *Shigella* species and *E. coli* genomes. When aligned to *S. flexneri*, *S. dysentariae*, and *S. boydii* sequences, none of the Lineage II or III specific alleles were 100% identical (Table 3). Furthermore, the other *Shigella* species did not house any lineage specific SNPs either, demonstrating the specificity of these SNPs to *S. sonnei*. Similarly, these SNPs and allele sequences also differentiated *S. sonnei* from *E. coli*, including the phenotypically similar enteroinvasive *E. coli*. SRR5943575 and SRR5943576 were predicted as not *S. sonnei* based on the results of the prediction, so SerotypeFinder was used to identify the serotype of these strains (30). Interestingly, they were both serotyped as O7:H18, which is not a possible serotype of *S. sonnei*. The serotype and lineage prediction results indicated that these isolates may have been mislabeled *E. coli* or misidentified as *Shigella*.

**Table 3:**
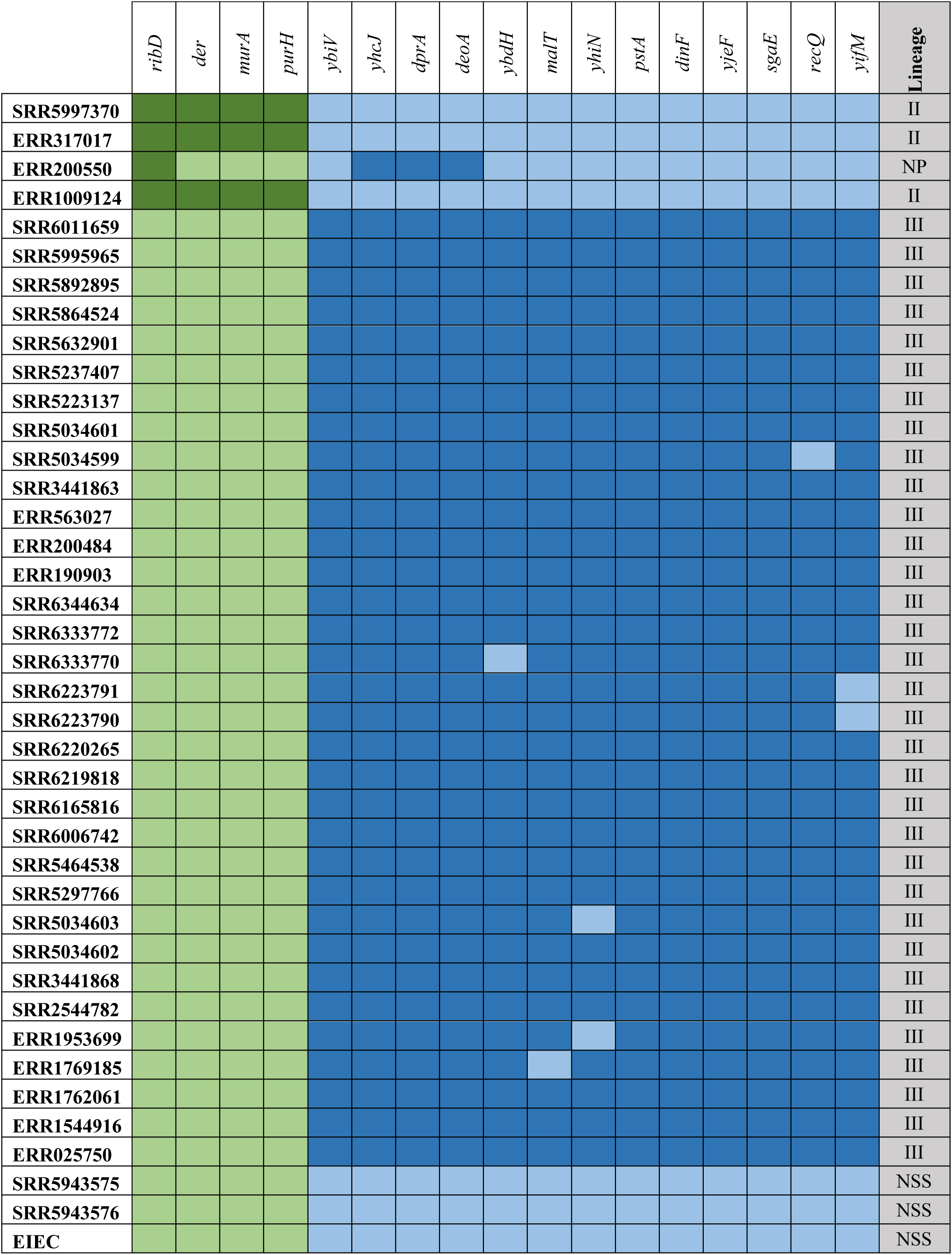
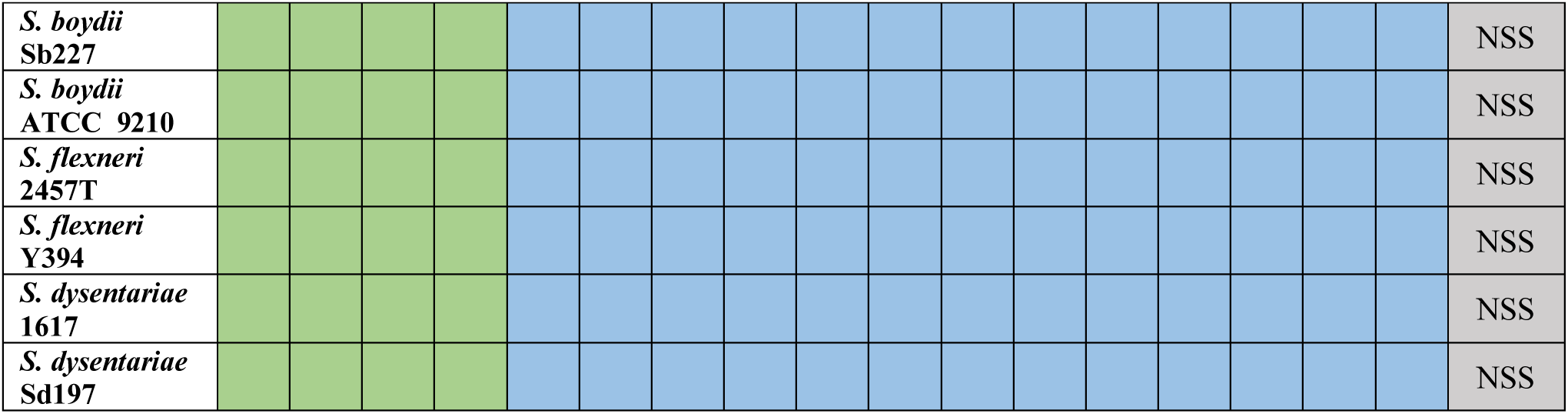
Global Lineage Prediction Tool: Nucleotide percent identity of the confirmed lineage specific alleles is represented by the coloring of the boxes. The dark green represents 100% nucleotide identity to Lineage II alleles and light green represents less than 100%. The dark blue represents 100% nucleotide identity to Lineage III alleles and light blue represents less than 100%. The grey-shaded column on the right lists the predicted Global Lineage for each strain. “NP” indicates non-predictable strains and “NSS” indicates isolates that were predicted as “Not *Shigella sonnei*.” The EIEC isolate is an enteroinvasive *E. coli* sequenced in a previous study by Pettengill et al. (35).

### Antimicrobial resistance of *S. sonnei* collection

Previous studies have shown that divergence of *S. sonnei* into the Global Lineages is partly due to the integration of AMR within the lineages (8). To investigate this interaction, the AMR genes within the *S. sonnei* genomes were identified (Fig. 4). All 22 of the *S. sonnei* isolates carried resistance genes for aminoglycosides and trimethoprim. Other resistance genes, like those for chloramphenicol resistance or macrolide resistance, were only found in 2 isolates each. Overall, the Lineage II isolates did not appear to carry more or less resistance genes than Lineage III, but there were some notable differences in the types of resistance they carried. The first is quinolone resistance: no Lineage II *S. sonnei* carried quinolone resistance determinants. Second, some Lineage II isolates carried a unique *bla*_TEM-1C_ gene for ampicillin resistance, while all Lineage III isolates that carried beta-lactam AMR genes had either the *bla*_TEM-1B_ or *bla*_OXA-1_ genes. Once the AMR genotype for each *S. sonnei* was determined, it was subsequently compared to the *in vitro* AMR profile provided from a previous analysis (Table 4) (16). Accordingly, genotypic resistance markers for macrolide, tetracycline, quinolone, chloramphenicol, beta-lactam, and sulfonamides aligned with the *in vitro* susceptibility testing performed previously. However, with AMR to trimethoprim and aminoglycosides, the genotype and phenotype did not correlate completely. Genome analysis found that all *S. sonnei* carried genes for trimethoprim resistance, but only 19 out of 22 isolates were classified as resistant by *in vitro* testing. Aminoglycoside resistance had a similar interaction, where all *S. sonnei* carried resistance genetic determinants but only 20 were found resistance. Though these genes were identified in the genomes of the *S. sonnei* isolates, these results indicate that there were discrepancies between phenotypic and genotypic resistance methods and that identification of resistance genes within a genome may not necessarily infer resistance.

**Figure 4:**
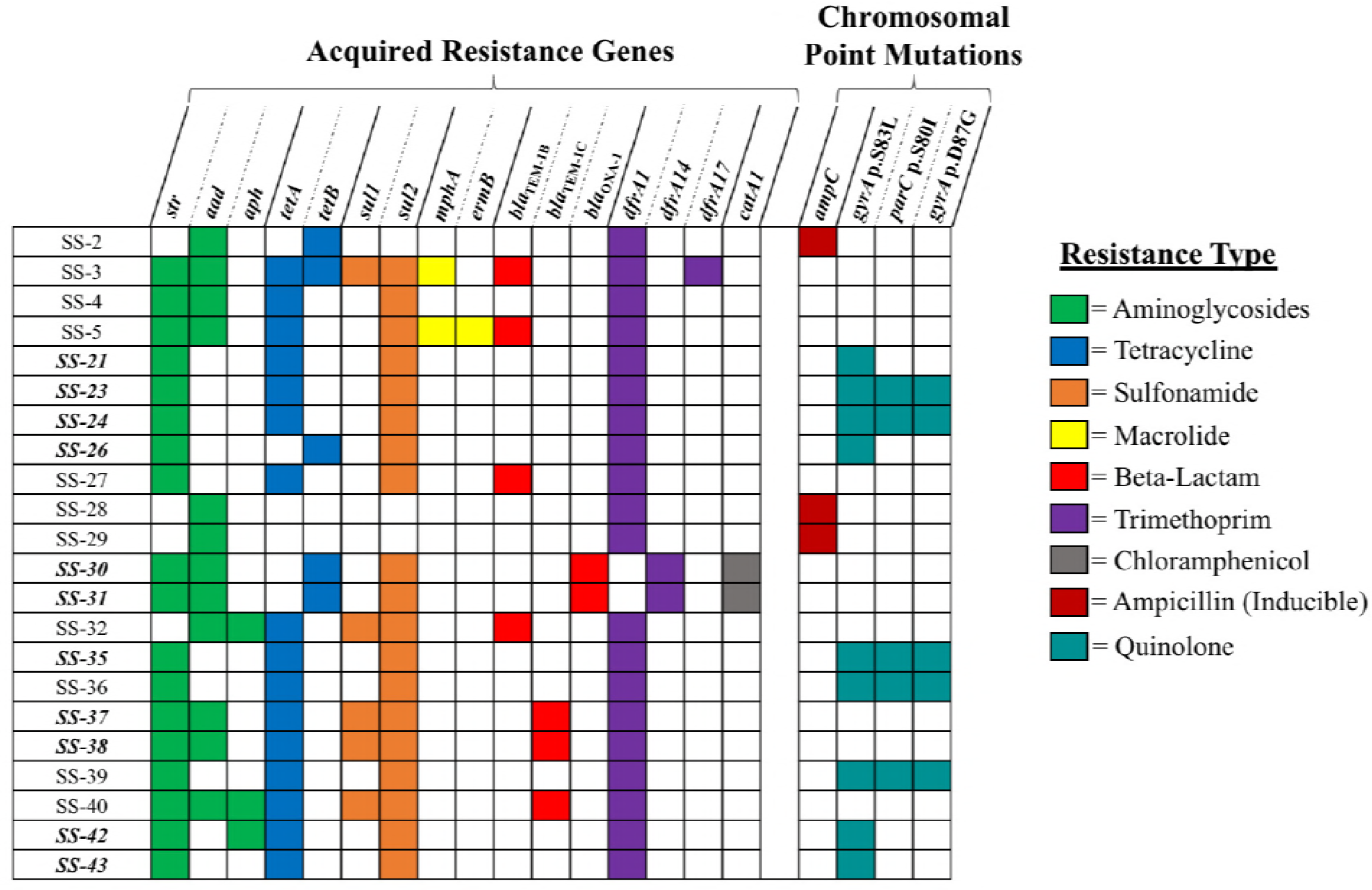
Antimicrobial Resistance Genes of *S. sonnei* Collection. This heat map is a representation of the different antimicrobial resistance genes identified in the *S. sonnei* sequences. The PSU-ID is listed on the left, followed by a list of genes read from left to right. The bolded and italicized PSU-IDs indicate international *S. sonnei*. If the square is colored, it indicates that the isolate carries the resistance gene with high identity and query coverage (>99%). The color of the square corresponds with the resistance type key on the right side. All genes shown were identified by both BARRGD and ResFinder 3.0, with the exception being the chromosomal point mutations, which were only identified by ResFinder 3.0.

**Table 4:**
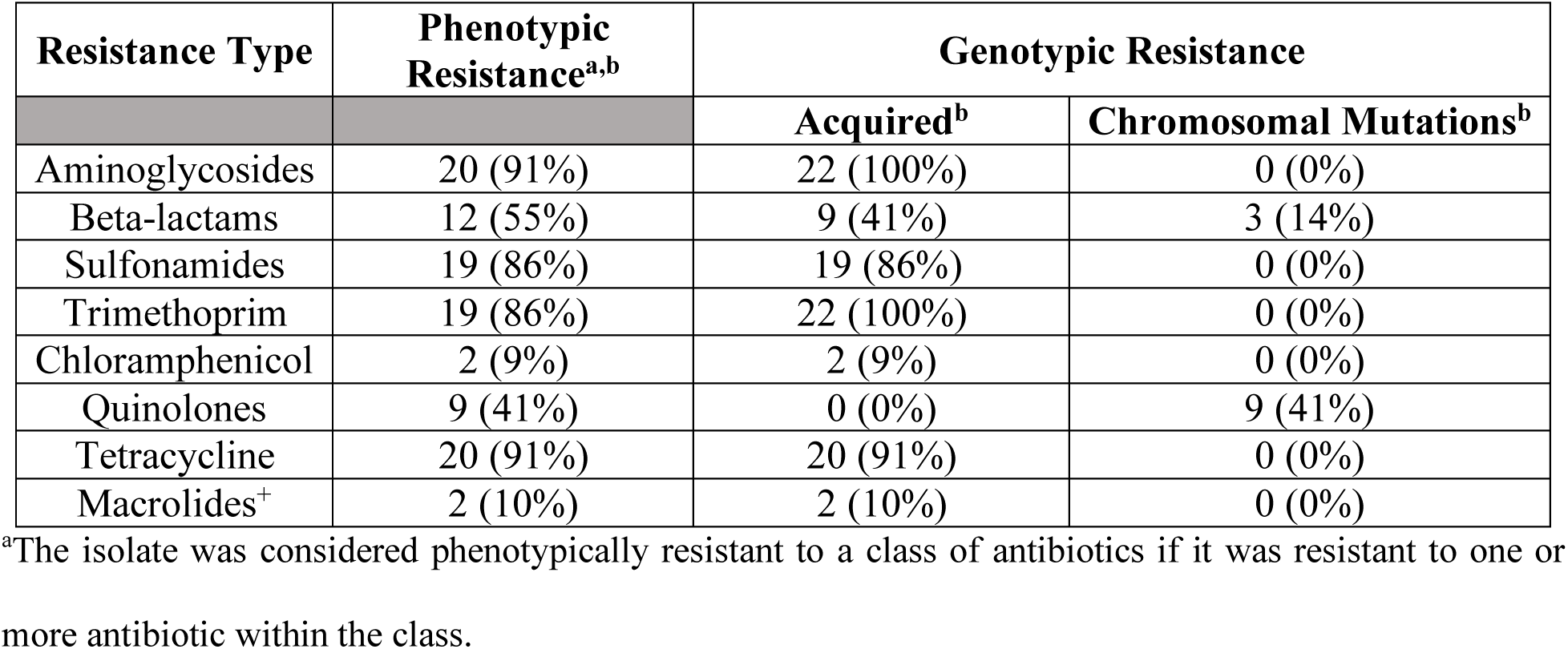
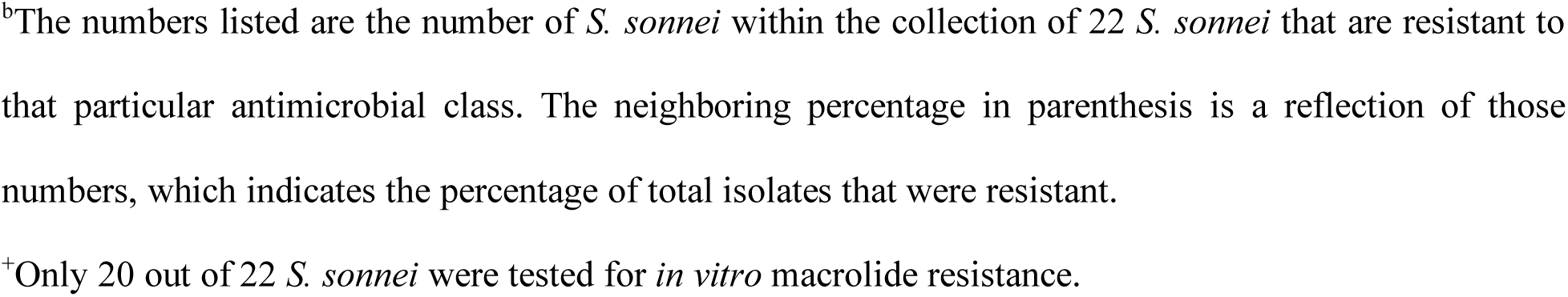
Summary of Antimicrobial Resistance Genotype and Phenotype Comparisons

| Resistance Type | Phenotypic Resistance<sup>a,b</sup> | Genotypic Resistance |  |
| --- | --- | --- | --- |
|  |  | Acquired<sup>b</sup> | Chromosomal Mutations<sup>b</sup> |
| Aminoglycosides | 20 (91%) | 22 (100%) | 0 (0%) |
| Beta-lactams | 12 (55%) | 9 (41%) | 3 (14%) |
| Sulfonamides | 19 (86%) | 19 (86%) | 0 (0%) |
| Trimethoprim | 19 (86%) | 22 (100%) | 0 (0%) |
| Chloramphenicol | 2 (9%) | 2 (9%) | 0 (0%) |
| Quinolones | 9 (41%) | 0 (0%) | 9 (41%) |
| Tetracycline | 20 (91%) | 20 (91%) | 0 (0%) |
| Macrolides<sup>+</sup> | 2 (10%) | 2 (10%) | 0 (0%) |

<sup>a</sup>The isolate was considered phenotypically resistant to a class of antibiotics if it was resistant to one or more antibiotic within the class.
<sup>b</sup>The numbers listed are the number of *S. sonnei* within the collection of 22 *S. sonnei* that are resistant to that particular antimicrobial class. The neighboring percentage in parenthesis is a reflection of those numbers, which indicates the percentage of total isolates that were resistant.
<sup>†</sup>Only 20 out of 22 *S. sonnei* were tested for *in vitro* macrolide resistance.

## Discussion

Previous studies of *S. sonnei* that have investigated phylogenetic segmentation align with the original analysis performed by Holt et al.; all *S*. sonnei appear to cluster into the Global Lineages (8, 9, 29, 31, 32). The original study, however, did not contain *Shigella* from the United States, so lineage characterization of isolates originating from the United States was not performed until Chung The et al. examined a collection of quinolone-resistant *S. sonnei* which included a few isolates from the United States (32). This study only utilized ciprofloxacin resistant specimens, but the researchers did find that the *S. sonnei* from the United States segmented into Lineage III. This aligns with the observations in this study; 18 of the isolates clustered into Lineage III (Fig. 2), and 9 of these isolates were resistant to fluoroquinolones (Fig. 4). Furthermore, Kozyreva et al. applied WGS to two separate *S. sonnei* outbreaks in California and found that all of the isolates from these outbreaks segmented into Lineage III (9). They also performed this phylogenetic analysis with some additional historic *S. sonnei* strains, in which some of the isolates collected prior to 2000 fell into Global Lineage 1 or 2. However, a similar temporal separation of Lineage II and III in the *S.sonnei* collection used in this study was not observed; though we did not include any isolates collected prior to 2009, isolate collection date did not appear to impact the lineage segmentation.

As WGS continues its integration into bacterial pathogen analysis, the development of tools for rapid identification and characterization is in the forefront of study. Tools that can identify specific gene sequences from either assembled genomes or raw sequencing reads to determine species, serotype, and other genetic factors have increased in demand due to the implementation of genomic sequencing methods. With this in mind, the lineage prediction allele sequences were determined to not only identify SNPs influencing the divergence of Lineage II and III, but they can also be utilized to predict phylogenetic characterization for timely isolate analysis. The 17 lineage specific alleles described here are original to this project, however one other group has examined the possibility of predicting lineages using SNP differences between *S. sonnei* isolates. Sangal et al. identified 4 gene variant sequences with lineage specific SNPs to predict lineages via high-resolution melting analysis (33); these alleles showed high sensitivity for identifying lineage segmentation, and one of the alleles they identified, *deoA*, was also identified by the methods used in this study. Adding to the findings of this previous research, we identified 16 additional alleles with lineage specific SNPs with a primary focus on differentiating Lineage II and III *S. sonnei* isolates. Lineage II and III prediction was of particular interest as both the findings in this study along with other previous reports find most *S. sonnei* isolates from both the United States and worldwide falling within either Lineage II or III. Also, these alleles showed high specificity for *S. sonnei* isolates; *E. coli* genomes and the genomes from other *Shigella* species, when aligned to the 17 lineage specific allele sequences, did not carry any of the alleles at the same level of nucleotide identity as the *S. sonnei* genomes. Overall, we believe these particular sequences are not only specific to Lineage II or III *S. sonnei* sequences, but they are also unique to *S. sonnei* and can be used to differentiate this organism from other similar bacteria.

When using additional *S. sonnei* sequences to test the lineage prediction alleles, *S. sonnei*III. ERR200550 was part of a large collection of *S. sonnei* from Latin America investigated by Baker et al. to examine via WGS, and coincidentally, this sequence was one of a number of genomes that segmented into the newly described Global Lineage V (31). This observation aligns isolate ERR200550 appeared to carry a unique pattern of Lineage II and III specific alleles and related SNPs (Table 3). After performing the phylogenetic analysis to depict lineages, this isolate branched independently from the others, and it did not segment into any known lineage, indicating that it may represent a new lineage or possibly an intermediately lineage between Lineage II and III. ERR200550 was part of a large collection of *S. sonnei* from Latin America investigated by Baker et al. to examine via WGS, and coincidentally, this sequence was one of a number of genomes that segmented into the newly described Global Lineage V (31). This observation aligns with the data presented in this work, as based on the lineage prediction test, this isolate carried a unique result and it appeared to be a hybrid strain between Lineage II and III. It is also an indication that, while the lineage prediction alleles were originally researched to predict Lineage II and III *S.sonnei*, the principles of the lineage prediction test could be modified to predict Lineage V sequences as well.

AMR in *S. sonnei* is important to examine not solely due to its human health implications, but also due to its possible influence on the evolution of *S. sonnei*. Acquisition of both acquired AMR genes and chromosomal mutations resulting in resistance have both been linked as a possible driver of *S. sonnei* phylogenetic divergence (8, 29) and the increase in prevalence of *S. sonnei* infections in developed regions over *S. flexneri* (6). To investigate this phenomenon and to determine the correlation of genetic resistance determinants to the previous *in vitro* susceptibility testing results within our collection of *S. sonnei*, known AMR determinants were identified within the genomes and then compared to phenotypic test results. As mentioned previously, there were differences in the types of AMR seen between the two lineages found in the collection; only Lineage III isolates carried resistance to quinolones and macrolides, the two drugs of major concern especially in the United States. This observation has been seen in previous literature, as Lineage III *S. sonnei* have been found to harbor more AMR genotypes when compared to other lineages (8, 9, 29, 31, 32). In regards to quinolone resistance, which has been a primary focus especially in international *S. sonnei* studies, Lineage III *S. sonnei* isolates appear to solely carry chromosomal mutations linked to quinolone resistance. Indeed, this observation was found not only in our collection, but also among other *S. sonnei* collections as well; some hypothesize that Lineage III *S. sonnei* are more likely to acquire both chromosomal mutations and genes that result in AMR due to selective pressures, regardless of geographic origin (29).

Investigations into the correlation of detecting AMR determinants from genome sequences and *in vitro* antimicrobial susceptibility testing have been performed on various pathogens with the purpose of determining if genomic resistance information can predict phenotype. In our *S. sonnei* collection, we saw correlation range from 86% to 100% based on the resistance detection methods we utilized; for beta-lactam, sulfonamide, chloramphenicol, quinolone, tetracycline, and macrolide resistance, we saw 100% correlation between genotype and phenotype, indicating that the genome would be a useful tool for predicting resistance in these cases (Table 4). Conversely, correlation in aminoglycoside and trimethoprim resistance was lower, approximately 91% and 86% respectively. In general, this data aligns with not only data from other similar bacterial pathogens like *Salmonella* and *E. coli* (12–15), but also with previous correlation results reported by Kozyreva et al. in *S. sonnei* (9). We did find differences in our work; we saw decreased correlation between genotype and phenotype in the aforementioned antibiotics, whereas Kozyreva et al. saw 100% correlation for these antimicrobials. It is difficult to discern the reason behind this discrepancy, as the antimicrobial susceptibility testing for these isolates was performed prior to this study. One possible explanation is breakpoint data for interpreting antimicrobial susceptibility testing results via MIC methodology may not align with genomic information. McDermott et al. described this interaction in nontyphoidal *Salmonella*, where they describe the breakpoint values of AST testing not correlating with genomic data for the aminoglycoside streptomycin (15). This could also be the case for *S. sonnei*; there are no CLSI-defined values for *Shigella* species in regards to aminoglycosides and trimethoprim as they are not utilized clinically (34), so the lack of correspondence between genetic resistance information and the *in vitro* testing results could be in response to non-representative MIC interpretation.

To the best of our knowledge, this is the first study to use WGS to examine *Shigella* isolates from Pennsylvania, and only the second to look specifically at *S. sonnei* isolates found in patients from the United States (9). We found that phylogenic analysis of *S. sonnei* adheres to the Global Lineage theory proposed by Holt et al., and that the segmentation of the strains into Lineage II and III can be predicted by a set of 17 allele sequences in the form of a lineage prediction tool. Additionally, we observed high correlation (greater than 85%) between the AMR genes and mutations and the *in vitro* antibiotic susceptibility testing data, along with finding differences in resistance between Lineage II and III isolates. Overall, we have contributed to both *S. sonnei* characterization and description using WGS.

## Acknowledgements

The authors wish to thank Lisa Dettinger, James Tait, and Yu Lung Li for their assistance with the preparation of the *S. sonnei* isolates. The authors also wish to thank Andrea Keefer for her assistance with sequencing quality control and Dr. Lingzi Xiaoli for her assistance with sequencing isolates. This work is supported by the U.S. Food and Drug Administration grant number 1U18FD006222-01 to Edward G. Dudley for support of GenomeTrakr in Pennsylvania, the Centers for Disease Control and Prevention Epidemiology and Laboratory Capacity (ELC) for collaboration in National Antimicrobial Resistance Monitoring System (CDC-RFA-CI10- 101204PPHF13), and the USDA National Institute of Food and Agriculture Federal Appropriations under Project PEN04522 and Accession number 0233376.

